# Inhibiting *Mycobacterium tuberculosis* ClpP1P2 by addressing the equatorial handle domain of ClpP1 subunit

**DOI:** 10.1101/713214

**Authors:** Yang Yang, Yibo Zhu, Tao Yang, Tao Li, Yuan Ju, Yingjie Song, Jun He, Huanxiang Liu, Rui Bao, Youfu Luo

## Abstract

Unlike other bacterial ClpP systems, mycobacterial ClpP1P2 complex is essential for mycobacterial survival. The functional details of *Mycobacterium tuberculosis* (*Mtb*) ClpP1P2 remains largely elusive and selectively targeting ClpP of different species is a big challenge. In this work, cediranib was demonstrated to significantly decrease the activity of *Mtb*ClpP1P2. By solving the crystal structure of cediranib-bound *Mtb*ClpP1P2, we found that cediranib dysregulates *Mtb*ClpP1P2 by interfering with handle domain of the equatorial region of *Mtb*ClpP1, indicating that the inter-ring dynamics are crucial for its function. This finding provides direct evidence for the notion that a conformational switch in the equatorial handle domain is essential for ClpP activity. We also present biochemical data to interpret the distinct interaction pattern and inhibitory properties of cediranib toward *Mtb*ClpP1P2. These results suggest that the variable handle domain region is responsible for the species-selectivity of cediranib, which suggests the equatorial handle domain as a potential region for screening pathogen-specific ClpP inhibitors.

## Introduction

Caseinolytic protease ClpP, a widely conserved self-compartmentalizing serine protease in bacteria, plays essential roles in protein metabolism and regulates diverse physiological functions including cell motility, genetic competence, cell differentiation, sporulation, and virulence, and is thus an attractive target for antibiotic development[1–8]. Notably, ClpP is an novel drug target for which both inhibition and activation result in an attenuated or lethal phenotype in many pathogens; thus agonists and antagonists aimed at the ClpP system represent promising drug candidates for further evaluation[9, 10]. The ClpP enzyme of *Mycobacterium tuberculosis (MTB), Mtb*ClpP1P2, exerts its proteolytic function by a heterotetramer of two protein subunits, ClpP1 and ClpP2[11, 12]. Moreover, encoding genes of subunits ClpP1 and ClpP2, have been shown to be essential genes for *MTB* survival and the deletion of either gene causes bacterial death[13]. A panel of *Mtb*ClpP1P2 inhibitors have been reported and they can be classified into two categories depending on their action modes. The first kind of small molecules act on the catalytically active center, covalently modifying the serine residues of the active sites of two subunits of *Mtb*ClpP1P2, including bortezomib, boron-containing analogues and beta lactones[14–17]. The second kind of inhibitors act on the chaperone protein (ClpX/ClpC1) binding site that competitively binds to the surface of the ClpP2 subunit. Representative molecules of this category are ADEP and its analogs[12]. Since the chaperones binding L/IGF region of ClpP2 subunit is highly conserved, as with the inhibitor of the *Mtb*ClpP1P2 catalytic center[18], the ADEP compounds are nonselective as well. As we know, the off-target effects may bring severe side effects during drug development. Thus it is desirable to find novel binding pocket or selective inhibitors targeting *Mtb*ClpP1P2 to avoid the potential adverse effects of nonselective ones. In this work, we demonstrate that cediranib, an orally available vascular endothelial growth factor receptor 2 (VEGFR-2) inhibitor, is able to inhibit *Mtb*ClpP1P2 peptidase activity. We solved the crystal structure of *Mtb*ClpP1P2 in complex with an agonist peptide and cediranib, and present details of the inhibitory mechanism. Unlike other ClpP inhibitors, cediranib dysregulates *Mtb*ClpP1P2 by interfering with the equatorial region of *Mtb*ClpP1, supporting the idea that the inter-ring dynamics are crucial for ClpP function. We also present biochemical data suggesting that this mechanism is distinct from those of previously reported ClpP inhibitors. These results reveal a novel druggable pocket in *Mtb*ClpP1P2, with potential implications for further research on ClpP catalytic mechanism and drug development.

## Results

### Identification of cediranib as a novel inhibitor of *Mtb*ClpP1P2

We screened about 2600 bioactive compounds (MedChemExpress, HY-L001) based on a peptidase activity assay. Eight compounds (hit rate 0.31%) that generated ≥80% inhibition of *Mtb*ClpP1P2 were selected for IC_50_ value evaluation (Fig 1A). Among these hits, cediranib and brivanib, which possesses an indolyl group, showed promising inhibitory effects toward *Mtb*ClpP1P2 peptidase activity; cediranib had a lower IC_50_ value (3.4 μM) than brivanib (12.5 μM) (Fig 1A and 1B). In addition, cediranib (100μM) led to a peak shift of 10°C in the thermal stability of *Mtb*ClpP1P2 in differential scanning calorimetry (DSC) assay, which confirms their direct interaction in solution (S1 Fig).

**Figure 1.**
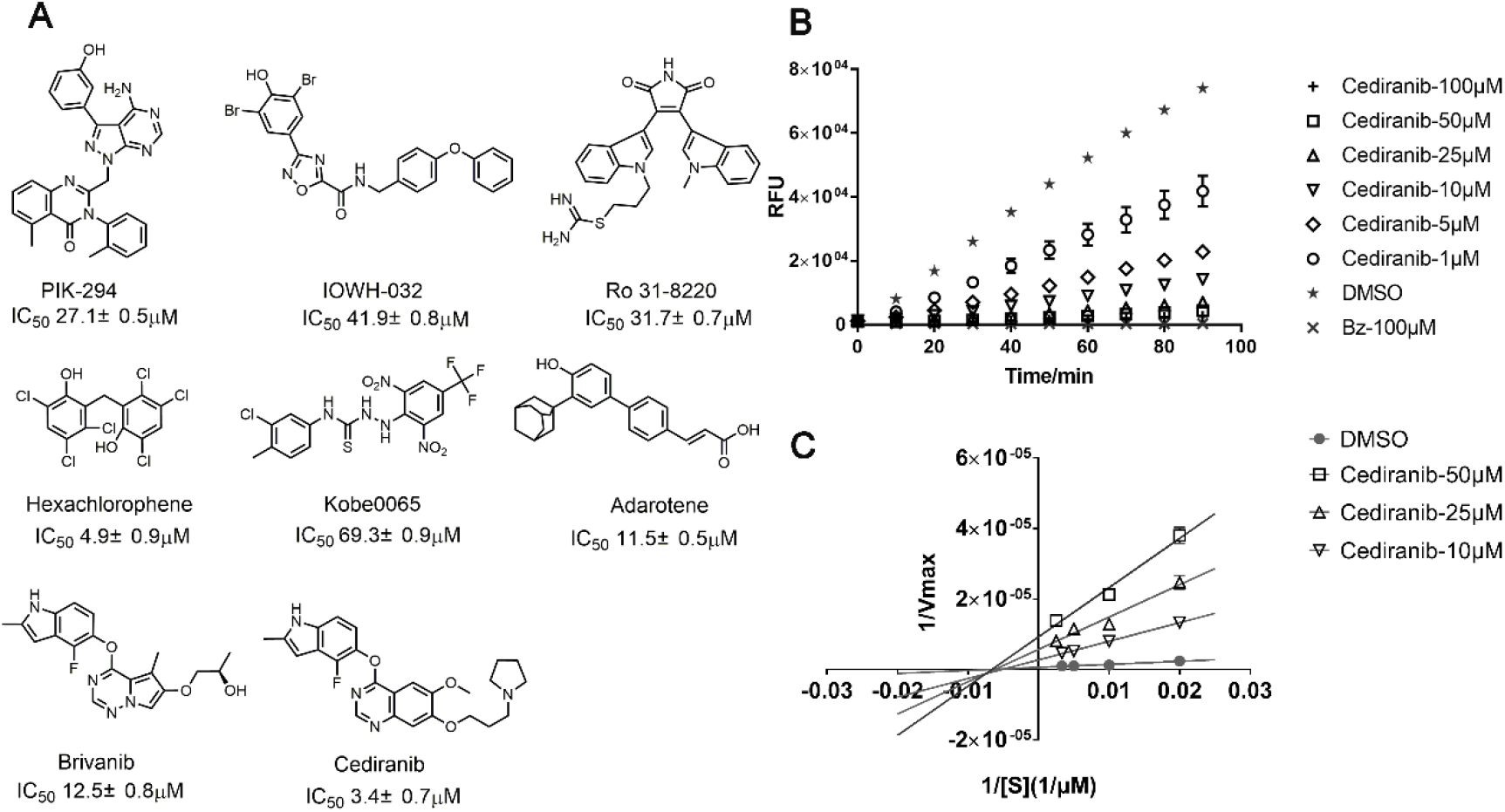
Cediranib inhibits *Mycobacterium tuberculosis* (*Mtb*) caseinolytic protease P1P2 (*Mtb*ClpP1P2) activity. (A) Chemical structure of compounds for secondary screening. (B) Cediranib inhibits *Mtb*ClpP1P2 cleavage of substrate Z-Gly-Gly-Leu-AMC in a peptidase assay. Concentrations of *Mtb*ClpP1P2 and Z-Gly-Gly-Leu-AMC were 0.5 μM and 100 μM, respectively. (C) Noncompetitive inhibition of *Mtb*ClpP1P2 cleavage of Z-Gly-Gly-Leu-AMC by cediranib. Cediranib increased the maximum reaction velocity (Vmax), but did not affect the Michaelis constant (Km) of the *Mtb*ClpP1P2 reaction.

To dissect whether cediranib is a species-selective inhibitor, we profiled its peptidase inhibitory effects against a panel of ClpPs from *Escherichia coli* (*Ec*ClpP), *Pseudomonas aeruginosa* (*Pa*ClpP1), and *Staphylococcus aureus* (*Sa*ClpP) (Table 1 and S2 Fig). The results show that cediranib has no obvious inhibitory effect on those ClpP variants. In contrast, cediranib demonstrated non-competitive inhibition of *Mtb*ClpP1P2 (Fig 1C), indicating that it may not form covalent bonds with the protease.

**Table 1.**
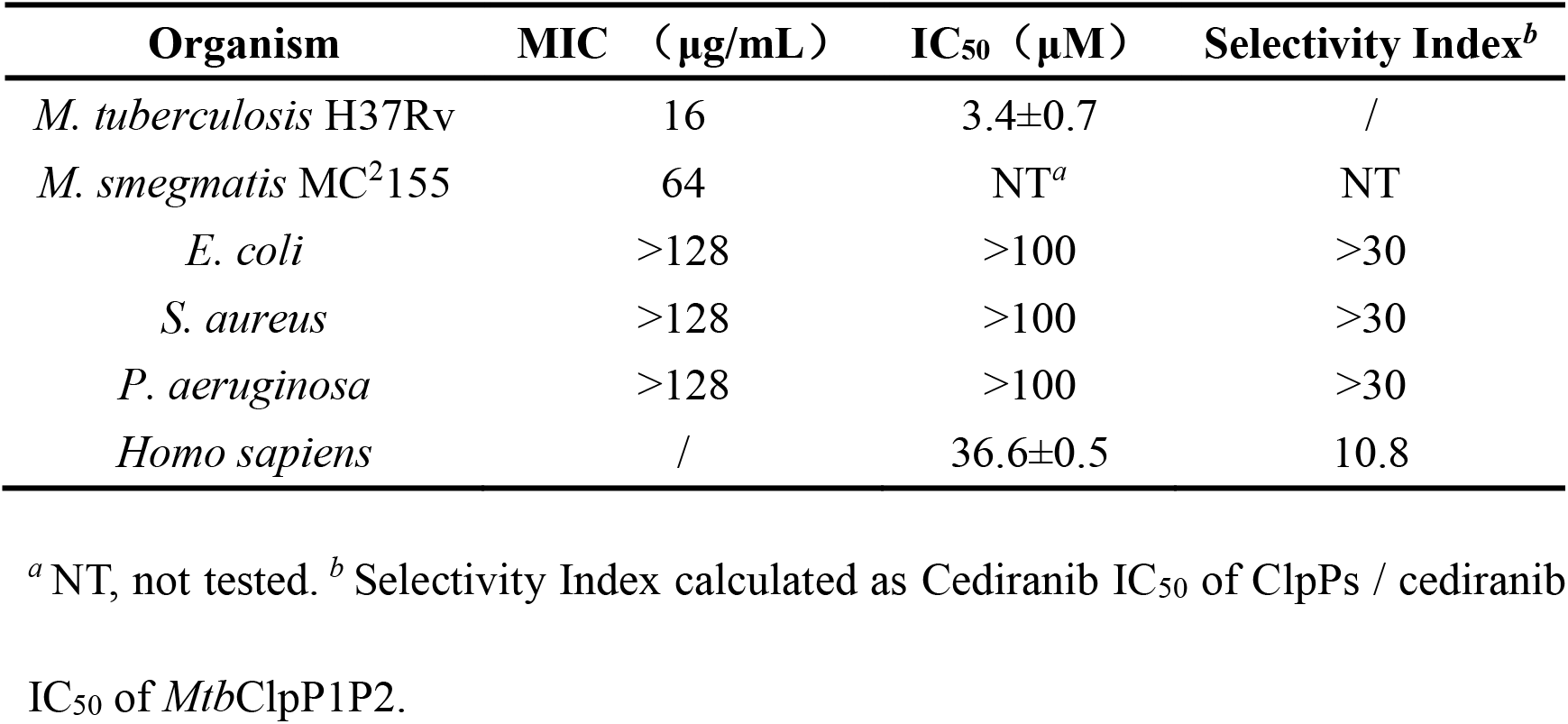
Minimum inhibitory concentrations (MIC) against different bacterial strains and the inhibition of ClpP peptidase activity by cediranib.

To validate the antibacterial activity of cediranib, we carried out growth inhibition assays on several pathogenic bacteria. As Table 1 and Table 2 show, cediranib had no inhibitory activity toward *S. aureus*, *P. aeruginosa*, *Escherichia coli* and *Enterococcus faecalis* (minimum inhibitory concentration [MIC] >128 μg/mL), while it was toxic to *E. faecium*, *Staphylococcus epidermidis, Klebsiella pneumoniae* and *Mycobacterium smegmatis* MC^2^155 (MIC 64 μg/mL). Cediranib inhibited the growth of *Mtb* H37Rv with an MIC value of 16 μg/mL. Therefore, cediranib exhibits species-selectivity in suppressing pathogen growth. Next, we performed peptidase assay on human ClpP (*h*ClpP) to estimate the toxicity of cediranib to human mitochondria. As shown in Table 1 and S2 Fig, cediranib displayed weak inhibition of *h*ClpP activity compared to *Mtb*ClpP1P2, with a selectivity index >10.

**Table 2.**
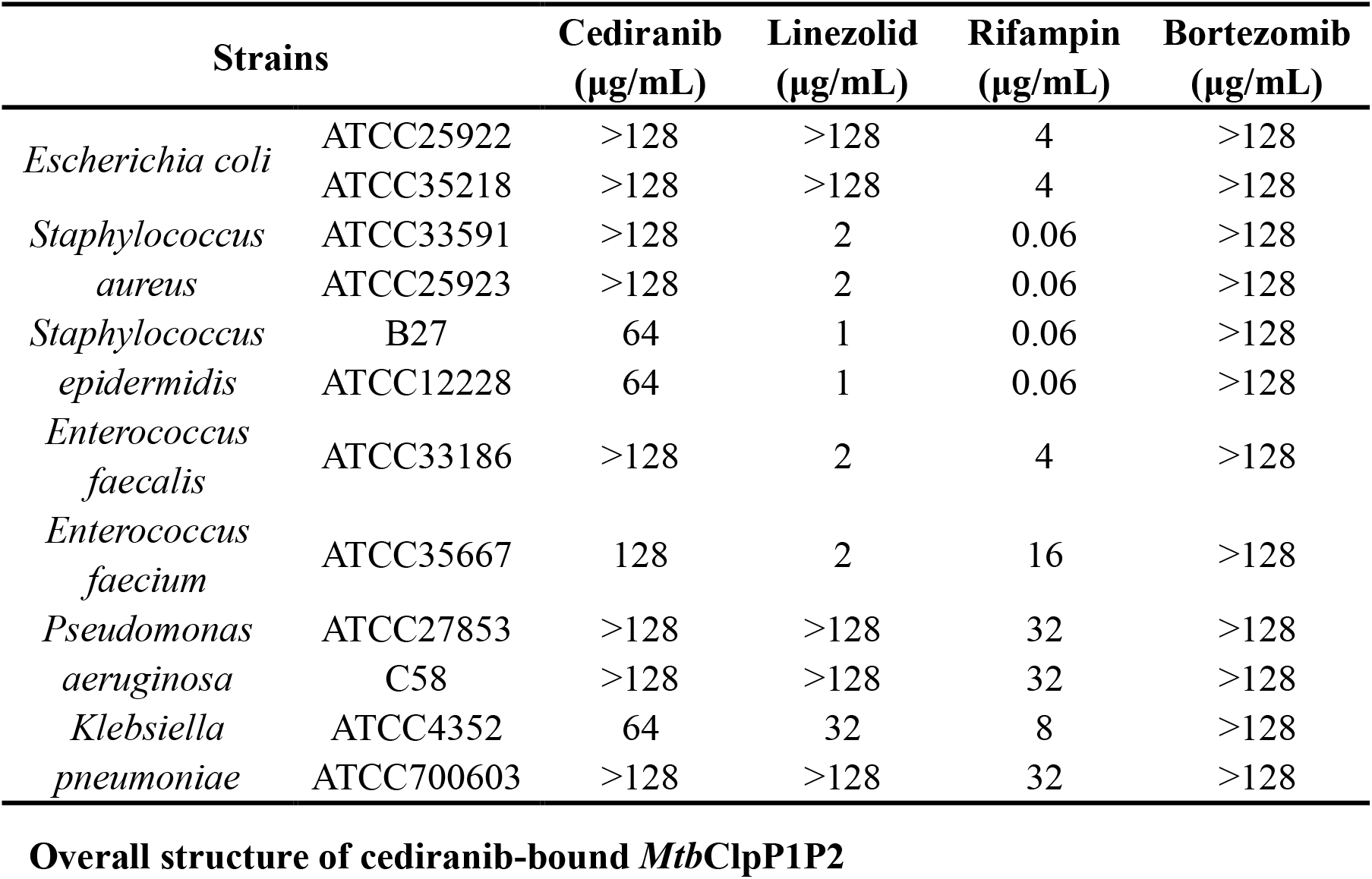
Minimum inhibitory concentration of cediranib against different strains.

To gain insights into the inhibitory mechanism of cediranib, we purified and crystallized *Mtb*ClpP1P2 in complex with cediranib and benzoyl-Leu-Leu (Bz-LL). The crystals belonged to space group C121 and diffracted to 2.7 Å resolution. The structure was determined by molecular replacement using the previously solved *Mtb*ClpP1P2 structure (PDB code 5DZK) as a search model, and the final model was refined to an Rfactor of 20.02% (Rfree =24.33%) (Table 3)[19]. The asymmetric unit has one hetero-tetradecameric complex structure composed of heptameric ClpP1 and ClpP2 rings (Fig 2A). The organization and overall structure remain identical to those in benzyloxycarbonyl-Ile-Leu (Z-IL)-bound and Bz-LL-bound *Mtb*ClpP1P2 structures[12, 19], presenting an open-pore conformation with 30 and 25 Å diameters in the ClpP1 and P2 rings, respectively (Fig 2B).

**Table 3.** Data collection and refinement statistics.

| Parameters | *Mtb*ClpP1P2 Cediranib Bz-LL |
| --- | --- |
| **Data collection** |  |
| Space group | *C121* |
| a, b, c(Å) | 210.63 180.78 95.93 |
| $\alpha$, $\beta$, $\gamma$(°) | 90.00 95.72 90.00 |
| Wavelength(Å) | 0.97915 |
| Resolution(Å) | 40.00 - 2.69(2.69 - 2.60) |
| CC<sub>1/2</sub> | 0.986(0.376) |
| Unique reflections | 96900(9595) |
| $R_{\text{meas}}(\%)^a$ | 20.8(134) |
| Mean $I/\sigma(I)^a$ | 8.04(1.04) |
| Completeness(%)<sup>a</sup> | 98.6(98.2) |
| Redundancy | 4.9(4.1) |
| **Refinement** |  |
| Resolution(Å) | 39.84 – 2.69 |
| $R_{\text{work}}/R_{\text{free}}^b$ | 0.2002/0.2433 |
| No.atoms |  |
| Protein | 19747 |
| Cediranib | 413 |
| Z-LL | 350 |
| Water | 187 |
| β-factors | |
| Protein | 49.86 |
| Cediranib | 68.36 |
| Z-LL | 55.26 |
| Water | 48.67 |
| R.m.s. deviations |  |
| Bond lengths(Å) | 0.009 |
| Bond angles(°) | 1.097 |
| Ramachandran plot |  |
| Favored (%) | 97.08 |
| Allowed (%) | 2.73 |
| Outliers (%) | 0.19 |

<sup>a</sup> The values in parentheses are for the outermost shell.
<sup>b</sup> $R_{\text{free}}$ is the $R_{\text{work}}$ based on 5% of the data excluded from the refinement.
$R_{\text{meas}} = \sum_{hkl} \sqrt{n/(n-1)} \sum_{i=1}^n |I_i(hkl) - \langle I(hkl) \rangle| / \sum_{hkl} \sum_i I_i(hkl)$ , where $\langle I(hkl) \rangle$ is the mean intensity of a set of equivalent reflections.
$R_{\text{work}} = \sum_{hkl} ||F_{\text{obs}}| - |F_{\text{calc}}|| / \sum_{hkl} |F_{\text{obs}}|$ , where $F_{\text{obs}}$ and $F_{\text{calc}}$ are observed and calculated structure factors, respectively.

**Figure 2.**
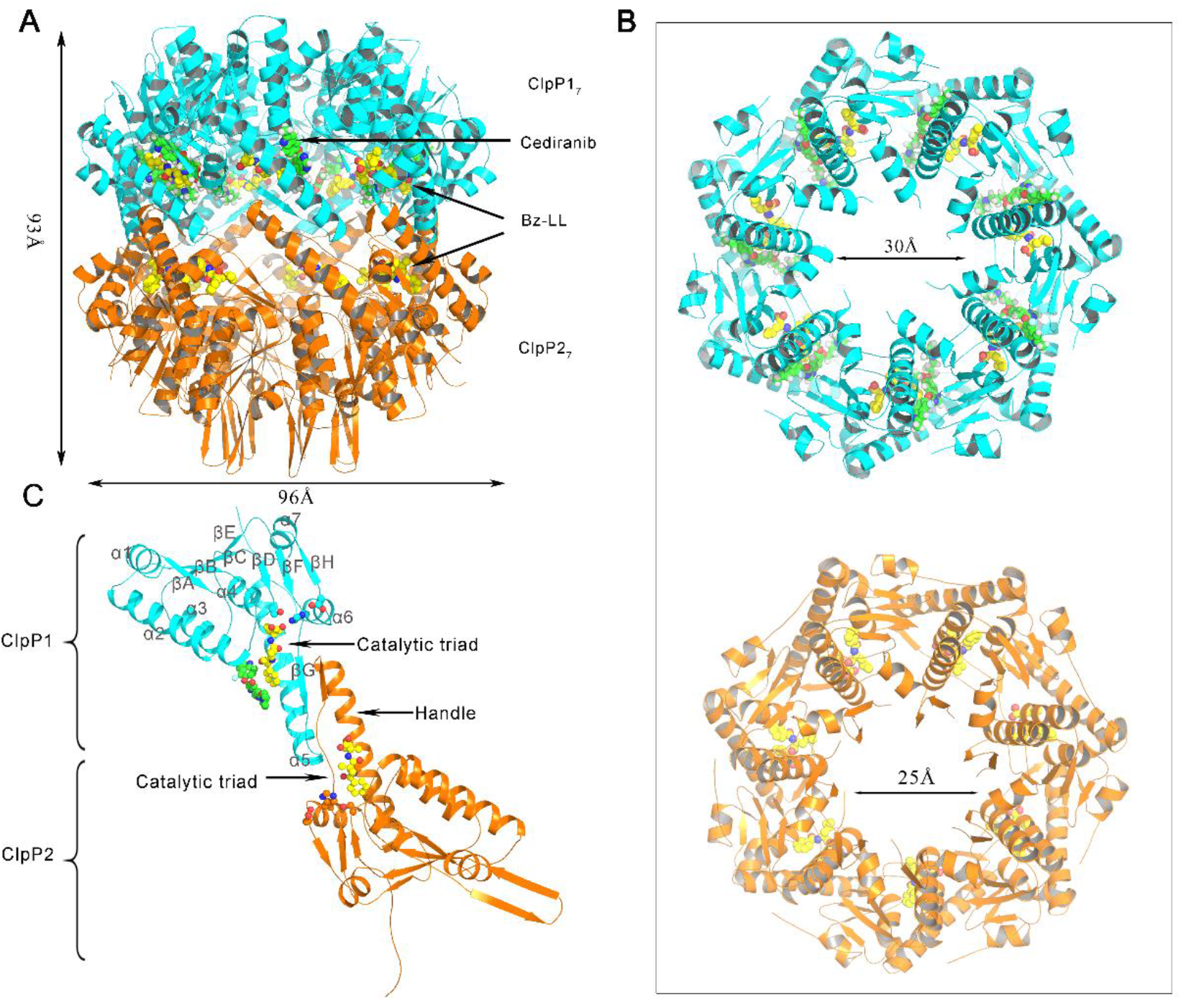
Cediranib-bound ClpP1P2 structures. (A) Side view of the *Mtb*ClpP1P2 tetradecamer. The *Mtb*ClpP1 heptamer (cyan subunits) and *Mtb*ClpP2 heptamer (orange subunits) are shown in cartoon representation. Spheres represent cediranib molecules (green) bound to the *Mtb*ClpP1 ring and Bz-Leu-Leu (Bz-LL) peptide (yellow) bound to the active sites of both rings. (B) Top panel, axial view of the *Mtb*ClpP1P2 tetradecamer from the *Mtb*ClpP1 side; bottom panel, axial view of the *Mtb*ClpP1P2 tetradecamer from the *Mtb*ClpP2 side. (C) *Mtb*ClpP1P2 monomer magnified from cediranib-bound *Mtb*ClpP1P2 structure. Secondary structure elements are labeled. The *Mtb*ClpP1P2 handle region and the catalytic triad residues are labeled. Cediranib and Bz-LL are colored green and yellow, respectively.

*Mtb*ClpP1 and *Mtb*ClpP2 share high sequence identity (48%) and have a common core structure, which adopts an α/β-fold consisting of two central twisted β-sheets flanked by seven α-helices and an extended handle domain (Fig 2C). The N-terminal regions of *Mtb*ClpP2 are in an extended β-hairpin conformation, and are further stabilized by the *Mtb*ClpP2 ring from the crystallographic symmetry unit. The typical catalytic triad (Ser-His-Asp) locates in the cleft between the core domain and the handle domain; all 14 active sites distribute on the inner surface of the assembled *Mtb*ClpP1P2 complex and eventually form the hydrolytic chamber. Bz-LL peptides occupied all cleavage centers in the *Mtb*ClpP1ClpP2 tetradecamer, whereas seven cediranib molecules bound within the cleft between *Mtb*ClpP1 monomers.

Self-association is a common and well-conserved property of Clp protease. The inter-ring interactions are mediated by the handle domain, a strand-turn-helix motif that forms the equatorial regions of the ClpP barrel[20–24]. Compared with the Bz-LL-bound *Mtb*Clp1Clp2 structure[19], the additional binding of cediranib slightly reduced the interface area between *Mtb*ClpP1 monomers (from 1356 Å2 to 1116 Å2), but does not change the assembly state of the whole complex, suggesting that the inhibitory mechanism of cediranib is different from that of compounds that disrupt the oligomerization of the ClpP complex[25].

### Cediranib-bound *Mtb*ClpP1P2 structure reveals a novel inhibitor binding-site

As previously reported[12, 19], the binding sites of Bz-LL and Z-IL are aligned parallel to β-strands 6 and 8 in each subunit and the agonist peptides form several hydrogen bonds with protein main-chain atoms. However, because *Mtb*ClpP2 has a longer loop after the βG strand (residues 127–129) and thus generates a shallower S1 pocket, Bz-LL is bound in opposite orientations in *Mtb*ClpP1 and *Mtb*ClpP2 (Fig 3A), supporting the notion that *Mtb*ClpP1 and *Mtb*ClpP2 have different substrate specificities[26]. In *Mtb*ClpP1, around the Bz-LL binding site, the averaged 2Fo–Fc electron-density map allowed us to unambiguously build the cediranib molecules between α5 and α3 (Fig 3B and 3C). Cediranib binds this site predominantly via hydrophobic interactions, while its indolyl group forms a cation–π interaction with Arg119 (Fig 2D). Additionally, the quinazoline ring of cediranib interacts with the aromatic rings from Trp174 and the benzoyl group of Bz-LL, resembling a π–π stacking interaction. In contrast to *Mtb*ClpP1, the corresponding sites in *Mtb*ClpP2 generate a more compact pore, where α5 moves closer to α3 and the large side chains of Met160 and Phe83 restrict accessibility to this pocket, making cediranib unable to bind to *Mtb*ClpP2 (Fig 3D).

**Fig 3.**
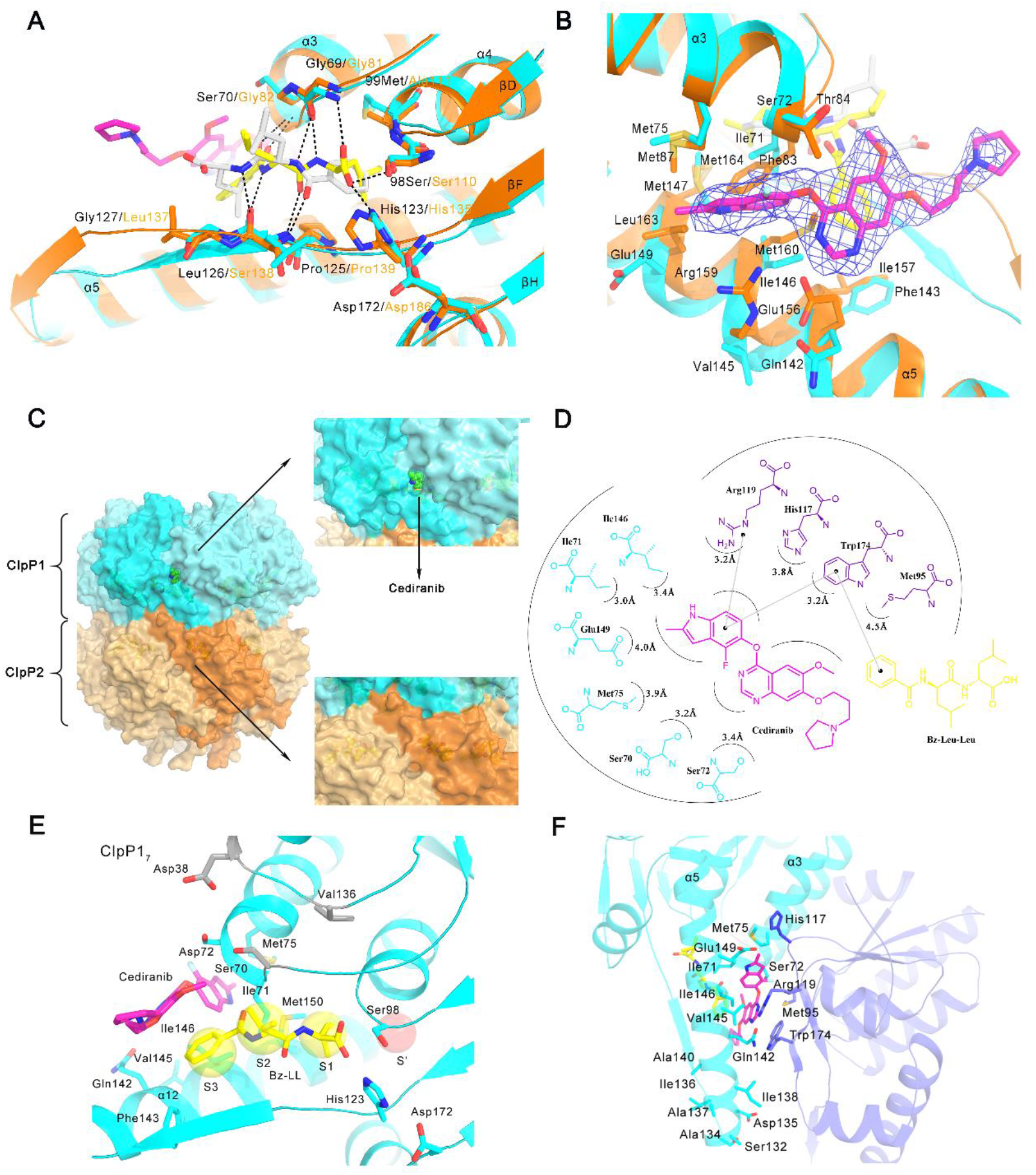
Structural details of cediranib binding pocket in *Mtb*ClpP1. (A) The activator is bound in opposite orientations in *Mtb*ClpP1 and *Mtb*ClpP2. Bz-LL in *Mtb*ClpP1 is colored yellow and Bz-LL in *Mtb*ClpP2 is colored gray. (B) 2Fo-Fc electron-density map of cediranib in *Mtb*ClpP1. (C) Surface representation of the *Mtb*ClpP1P2 tetradecamer with bound cediranib reveals that cediranib binds in a novel site. (D) A cartoon representation of the cediranib interactions with *Mtb*ClpP1. (E) Molecular interactions between cediranib and residues in the binding pocket. Cediranib does not interact directly with the active site, but interferes with the presumably defined equatorial pores and dynamic handle regions. (F) Detailed view of the cediranib-bound pocket formed by residues Gln142–Glu149 in α5, Ile71–Ala76 in α3, and three short β-sheets (βD, βF, βH) from the neighboring *Mtb*ClpP1 subunit.

Unlike the peptidyl inhibitors that occupy the S1–S3 subsites of ClpP[27], cediranib does not directly interact with the active site, but interferes with the dynamic handle region, which was proposed to be responsible for product release[21, 28](Fig 3C and 3E). Though a small channel in the variable handle region of *Mtb*ClpP1 was observed in ADEP-*Mtb*ClpP1P2 complex in previous study[12], we firstly demonstrated this small channel could be occupied by a small molecule, herein cediranib, and be employed to realize *Mtb*ClpP selectivity. As Fig 3F illustrates, the cediranib-binding pocket is formed by α5 (residues Gln142–Glu149), α3 (residues Ile71–Ala76) and three short β-sheets (βD, βF, βH) from the neighboring ClpP1 subunit.It is worth noting that α5 is the major part of the handle domain.

### Molecular dynamics simulation studies revealed cediranib interferes the conformational changes of *Mtb*ClpP1

Previous molecular dynamics simulation studies on SaClpP revealed that α5 adopts a kinked conformation by breaking the helix at residue Lys145 (corresponding to Val145 in *Mtb*ClpP1)[24]. During the reaction cycle, the handle domain undergoes an unfolding/refolding process, allowing the whole cylindrical ClpP barrel to adopt extended, compact and compressed states[23, 24, 29–31]. In order to analysis the effect of cediranib on the dynamic transition of handle domain, we performed a 500-ns MD simulation on *Mtb*ClpP1 dimer (chain B and M) with/without cediranib (S3 Fig). The r.m.s. deviation (RMSD) values of Cα from the head domain and handle domain were monitored, the results indicated that handle domain underwent dramatic conformational changes during the simulation, whereas both the head domain and handle domian of B chain was more stable than that of M chain (S3A Fig and S3C Fig). Next, the secondary structure transformation of handle domain were calculated along the trajectory of simulation by DSSP[24] (S3B Fig and S3D Fig). The profile revealed that the N-terminal part of helix E (residues 133–158) underwent a helical unfolding/refolding process. In the absence of cediranib, the N-terminal part (residues 133–137) in chain B primarily adopted turn structure in 0-350 ns. Then, residues 133–154 adopted coil, turn and 3-10Helix structures. In chain M, residues 143–147 mainly adopted turn structure. With binding of cediranib, the N-terminal part(residues 133–137) of chain B adoped coil and turn strcture in the initial stage (0–300 ns). Subsequently(300-500ns), these residues gradually readopted some 3-10Helix and α-helix. By contrast to chain B, chain M was much more unstable, particularly at 250ns, most α-helix structure adopted turn and coil (S3B Fig and S3D Fig). To identify the most significant motions of handle domain of *Mtb*ClpP1 dimer induced by cediranib, we performed PCA using the MD trajectory[24]. As shown in S3E Fig and S3F Fig, during the conformational transition, four intermediate conformations (see the 50-ns, 100-ns,300-ns, and 500-ns snapshots in Fig 3E and 3F) was observed. Based on the results of MD simulation and PCA, we constructed a rough energy landscape for the conformational transition of *Mtb*ClpP1 dimer projected onto the first two principal components, PC1 and PC2. Obviously, the binding of cediranib to *Mtb*ClpP1 interferes the conformational changes, likely preventing the protease from accomplishing the enzymatic process.

### Mutagenesis and functional analysis of the cediranib-binding site

As we known, the ClpPs from different species share high sequence similarity (55.0–90.9%) and conserved overall structure (Fig 4A). Consistently, the general and critical gating mechanism of ClpP requires the handle domain to preserve its structural integrity[29]. In the dynamic region of the handle domain (Ser132–Val145 in *Mtb*ClpP1), several sites involved in structural stabilization have been identified to be important for ClpP function: Gln132/Glu135 in *Sa*ClpP (corresponding to Ser132/Asp135 in *Mtb*ClpP1, interacting with Arg171)[23]; Ile149/Ile151 in *Ec*ClpP (corresponding to Ile136/Ile138 in *Mtb*ClpP1)[21]; and Ala153 in *Streptococcus pneumoniae* ClpP (corresponding to Ala140 in *Mtb*ClpP1) (Fig 3F)[22]. Thus, the cediranib-bound *Mtb*ClpP1P2 structure provides additional evidence to highlight the critical role of the handle region.

**Fig 4.**
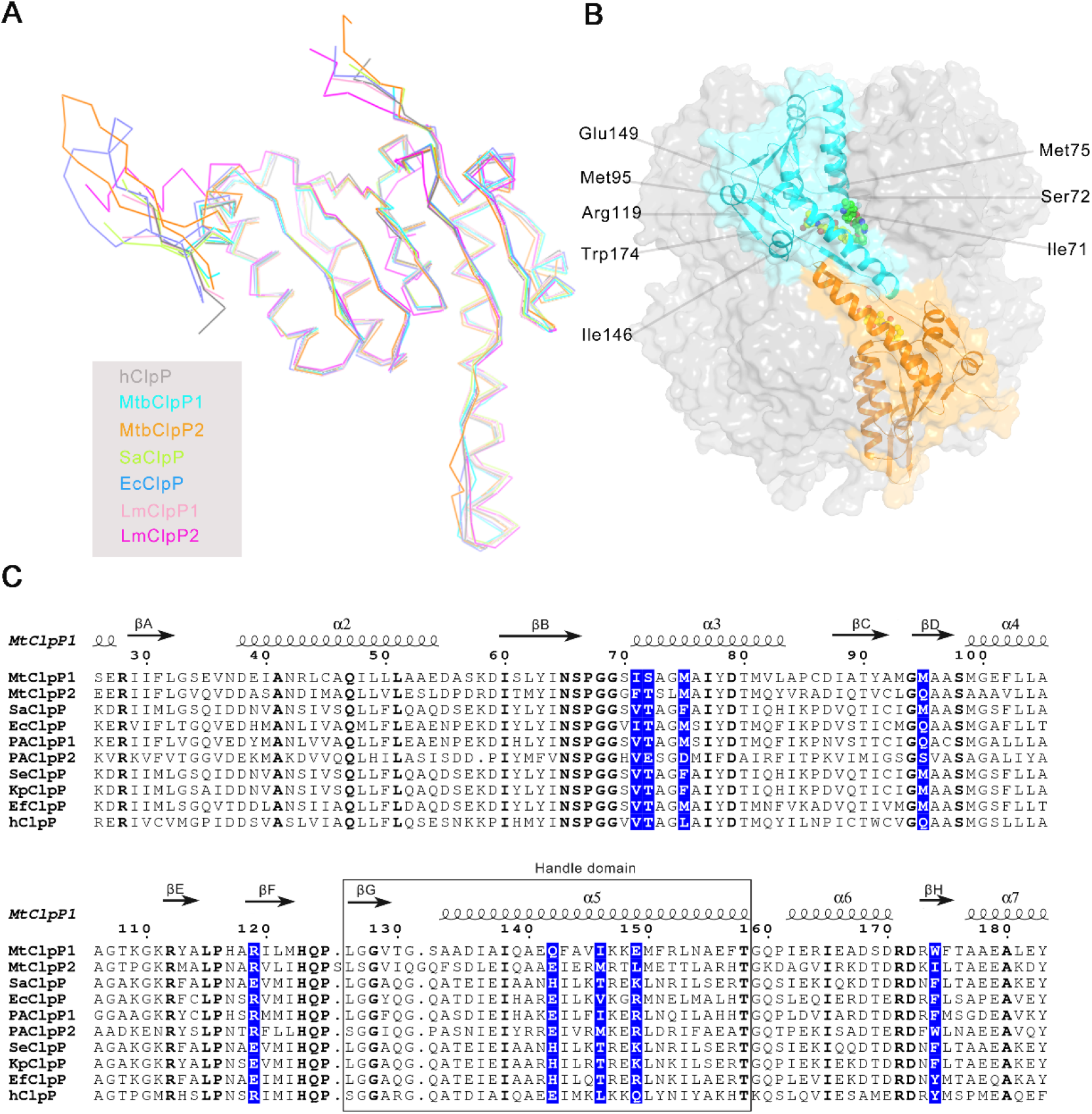
Importance of the handle domain and analysis of distinct ClpP sequences. Importance of the handle domain and analysis of distinct ClpP sequences. (A) Superposition and ribbon representation of the monomeric structures of *Mtb*ClpP1 (cyan), *Mtb*ClpP2 (orange), Staphylococcus aureus (Sa)ClpP (green), Escherichia coli (Ec)ClpP (light blue), Listeria monocytogenes (Lm)ClpP1 (pink), LmClpP2 (magenta), and human (h)ClpP (gray). (B) *Mtb*ClpP1P2 side views. Mutant residue sites are labeled. Spheres represent cediranib molecules (green) bound to the *Mtb*ClpP1 ring and Bz-Leu-Leu (Bz-LL) peptide (yellow) bound to the active sites of both rings. (C) Sequence alignment and secondary structure assignment of ClpPs. Sequence alignment was performed in BioEdit. Identical residues are highlighted in blue. Secondary structure elements present in the *Mtb*ClpP1P2 structure (PDB code 6IW7) are shown on the top of the sequence alignment.

To investigate the residues in the cediranib-binding pocket, we introduced different mutations in relevant sites based on sequence alignment (Fig 4B and Table 4). Consistent with the importance of the equatorial pore and handle domain, most site-directed mutations in the selected sites decreased or abolished *Mtb*ClpP1P2 activity. Considering Ile71, Met75, Ile146 and Trp174, even when residues with similar side chains were substituted in those sites, the enzyme activity was not retained, suggesting there are constraints on both residue size and chemical properties to maintain the local structural integrity and flexibility (Table 4 and Fig 4C).

**Table 4.**
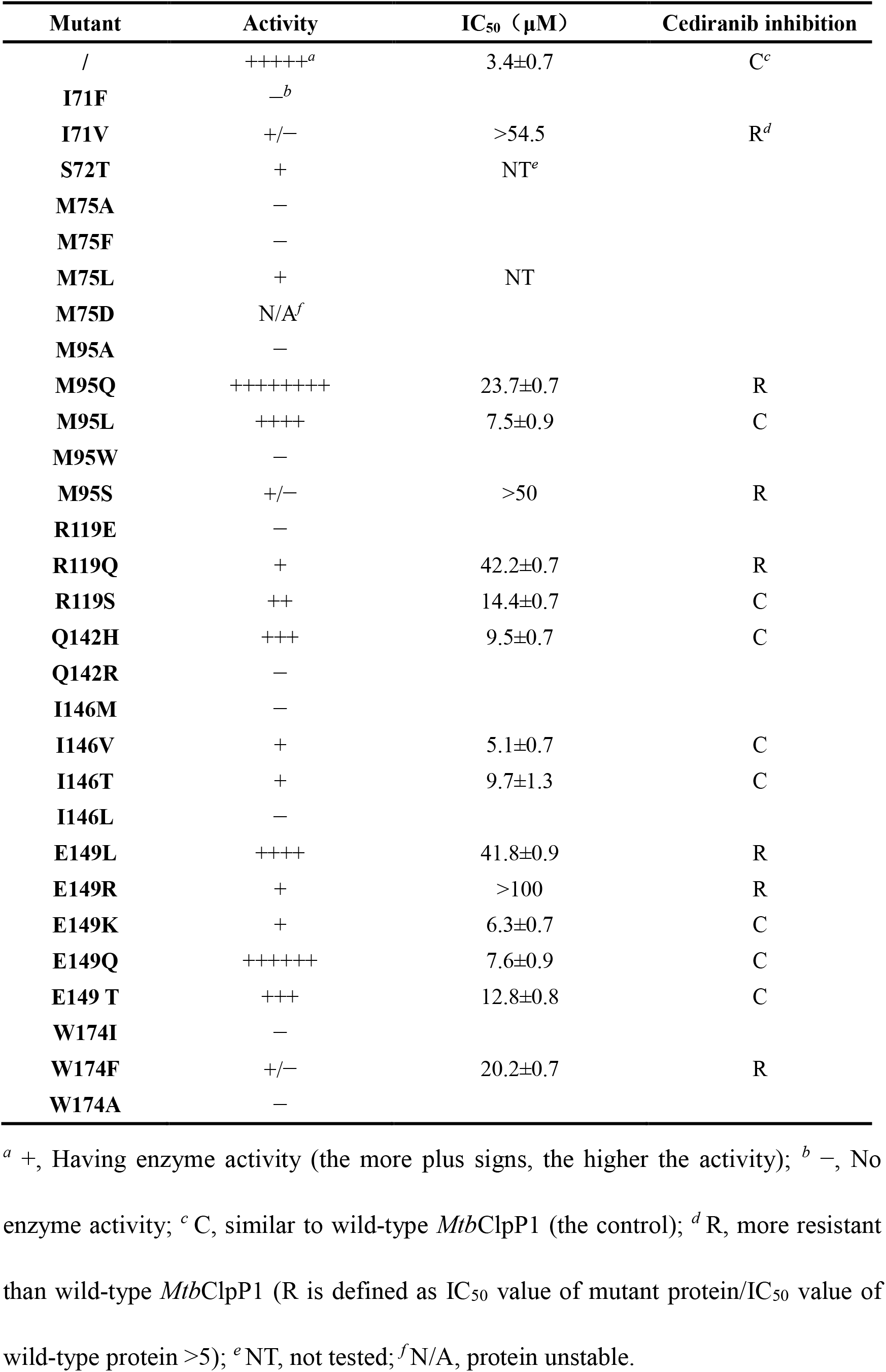
Mutagenesis analysis of *Mtb*ClpP1.

An electrostatic interaction between two *Mtb*ClpP1 subunits (Arg119–Gln142) (Fig 3F) seems to be an important factor mediating the side pore gating, because any substitution that changed the charge properties of either site resulted in reduced or abolished activity. In contrast, sites Met95 and Glu149 of *Mtb*ClpP1 exhibit various levels of tolerance to mutation. In particular, Met95Gln and Glu149Leu variants retain activity and have altered sensitivity of *Mtb*ClpP1P2 to cediranib, providing a structural basis for the specific recognition between cediranib and *Mtb*ClpP1 (Table 4).

## Discussion

ClpP represents a unique type of serine protease complex responsible for the proteolysis of damaged or misfolded proteins. Because it plays critical roles in the regulation of infectivity and virulence of many bacterial pathogens and is involved in various human diseases, interfering with ClpP activity has been evaluated as a potential therapeutic strategy in the treatment of different ailments[32]. In-depth understanding of the mechanism of ClpP reaction and regulation is crucial for development of chemotherapeutic agents that target this protein. The importance of the dynamic equatorial region in ClpP has long been recognized[21–24, 29–31].

As the first identified inhibitor targeting flexible handle region of ClpP, cediranib shows a distinct inhibitory mechanism. The cediranib-bound *Mtb*ClpP1P2 complex structure provides valuable evidence to support the important roles of equatorial handle domain for ClpP function^[21][26]^. Compared with the highly conserved critical residues in the N-terminal region of α5 (Ile136, Ile138, and Ala140 in *Mtb*ClpP1) (Fig 4C), residues participating in cediranib binding are relatively diverse among different ClpP homologs, leading to a substantial variation in the size of the binding pockets near to the side pores, which suggests this mechanism could be employed for pathogen-selective ClpP inhibitor development.

## Materials and Methods

### Bacterial strains and culture conditions

*Escherichia coli* DH5α and BL21 (DE3) were cultured at 37°C in Luria broth (LB) and LB-agar plates. DNA encoding *M. tuberculosis* ClpP1 and ClpP2 (spanning residues 7–200 and 13–214, respectively) was cloned into a pRSF-Duet vector preceded by an N-terminal His6-SUMO tag (Novagen)[11, 12]. Full-length *Ec*ClpP, *Sa*ClpP, *Pa*ClpP1 and the *h*ClpP gene without its mitochondrial targeting sequence (residues Met1–Pro57) were also respectively cloned into pRSF-Duet preceded by an N-terminal His6-SUMO tag[33–35]. PCR was performed with PrimeSTAR^®^ max DNA polymerase (Takara) and *Ex Taq*^®^ polymerase (Takara). The plasmids carrying distinct ClpP genes were transformed into *Escherichia coli* BL21 (DE3) cells (TransGen Biotech). The plasmids and primers used in this study are listed in Appendix Table S1 and Appendix Table S2.

### Protein expression and purification

*M. tuberculosis* ClpP1 (residues 7–200) and ClpP2 (residues 13–214) with N-terminal His_6_-SUMO tags were expressed and purified as described, with minor modifications[11, 12, 14]. In short, ClpP1 and ClpP2 were individually overexpressed in *E. coli* strain BL21 (DE3) in LB broth at 16°C, following induction with 0.5 mM isopropyl β-D-1-thiogalactopyranoside for about 16 h with shaking at 220 rpm in medium supplemented with 50 μg/mL kanamycin. Cells were resuspended and lysed in buffer containing 50 mM K_2_HPO_4_/KH_2_PO_4_, pH 7.6, 500 mM KCl, 10% glycerol and 0.5 mM dithiothreitol. His-tagged *Mtb*ClpP1/*Mtb*ClpP2 proteins were purified using a nickel affinity column and eluted with the same buffer supplemented with 250 mM imidazole. The His_6_-SUMO tag of eluted proteins was removed using ubiquitin-like-protease 1 (ULP1) at 4°C. The tag-free proteins were further purified by anion exchange chromatography (MonoQ; GE Healthcare). Peak fractions were collected, concentrated and applied to Superdex 200 16/600 (GE Healthcare) preequilibrated with buffer containing 10 mM HEPES, pH 7.5, 50 mM NaCl and 2 mM dithiothreitol for final purification. Purified ClpP1 and ClpP2 were concentrated in Amicon centrifugal concentration devices (Millipore) to >80 μM tetradecamer. *Ec*ClpP, *Sa*ClpP, *Pa*ClpP1 and *h*ClpP proteins were expressed as described above. These proteins were purified using a nickel affinity column and then incubated with ULP1 at 4°C to cleave the His6-SUMO tag[36].

### Compound screening

The peptidase activity of *Mtb*ClpP1P2 was monitored at 30°C in black 96-well plates, as described[11, 14]. After peptide bond cleavage, the fluorophore 7-amino-4-methylcoumarin of the fluorogenic substrate Z-Gly-Gly-Leu-AMC is liberated and can be quantified. Briefly, each well contained 100 μM fluorogenic peptide, 0.5 μM tetradecamer ClpP1P2, and 2 mM Bz-LL in 80 μL of buffer containing 50 mM phosphate buffer, pH 7.6, 300 mM KCl, 5 mM MgCl_2_ and 5% glycerol. Substrate cleavage was monitored using an all-in-one microplate reader (Gen5; BioTek) by exciting at 380 nm and following the increase in fluorescence emission at 460 nm. The deviation of fluorescence value in at least two independent measurements was ≤5%. The screening assay was carried out in 96-well format as described above, and about 2600 compounds were screened at 100 μM. Positive (bortezomib) and negative controls (dimethyl sulfoxide; DMSO) were included on every plate and were used to assess the performance of the primary screen. Bz-LL was purchased from GL Biochem (Shanghai) Ltd. Cediranib was purchased from MedChemExpress.

### IC_50_ determination

To determine the potency of the 8 “hit” compounds against *MtbClpP1P2,* these compounds were tested at concentrations ranging from 1 to 100 μM in a 96-well plate format. The reaction mix contained 0.5 μM *Mtb*ClpP1P2 tetradecamer and 2 mM Bz-LL, with 0.8 μL of each dilution of the compound or DMSO in a total volume of 80 μL. The reaction was initiated by addition of 0.8 μL of 10 mM fluorogenic substrate Z-Gly-Gly-Leu-AMC (100 μM final concentration) to the reaction mix. Initial velocity data was obtained by the monitoring increase in the fluorescence due to hydrolysis of the substrate using the microplate reader at 10-min intervals over 60 min. The IC_50_ value for each compound was obtained by nonlinear regression curve fitting of a four-parameter variable slope equation to the dose– response data using Prism software.

The peptidase activity of *Ec*ClpP, *Sa*ClpP, *Pa*ClpP1 and *h*ClpP was determined using a fluorescence-based assay with Suc-LY-AMC as the substrate, according to literature protocols[33–35]. The reaction mix contained 0.5 μM *Ec*ClpP/2.5 μM *Sa*ClpP/0.5 μM *Pa*ClpP/3 μM *h*ClpP tetradecamer, with 0.8 μL of each dilution of the test compound or DMSO in a total volume of 80 μL. The IC_50_ values for cediranib against *Ec*ClpP, *Sa*ClpP, *Pa*ClpP1 and *h*ClpP were obtained as described above.

Non-competitive inhibition was investigated by using 0.5 μM *Mtb*ClpP1P2 tetradecamer and 2 mM Bz-LL with different concentrations of cediranib. An equivalent volume of DMSO was added to the control group. The serially diluted substrate (500, 400, 300, 200, 100 and 50 μM) was added into wells of a 96-well plate and incubated for 30min at room temperature. Fluorescence was measured at 30°C using the microplate reader (excitation, 380 nm; emission, 460 nm) for 1 h[9].

### Crystallization

Cediranib-*Mtb*ClpP1P2 complex crystals were prepared by the hanging drop method referring to previously published works[12, 19]. The precipitant solution consisted of 1.5 M (NH_4_)_2_SO_4_ and 0.1 M MES, pH 6.5. A drop contained a mixture of 2 μL protein (about 2.5 mg/mL ClpP1 and ClpP2, 0.2 mM cediranib maleate, 5 mM Bz-LL, 10 mM HEPES, 50 mM NaCl, 2 mM DTT, pH 7.5) and 2 μL of precipitant solution and was incubated at 18°C for about 3 months. Crystals were soaked briefly in 2 M Li_2_SO_4_ solution and were stored in liquid nitrogen.

### Data collection and structure determination

The X-ray data were collected with a CCD camera at station BL-19U of the Shanghai synchrotron radiation facility (SSRF), Shanghai, China. The diffraction data were indexed, integrated, and scaled using the HKL2000 program suite[37]. The process of structure building and refinement was monitored using the COOT and PHENIX suites[38, 39]. The PDB code for the co-crystal structure of cediranib with *Mtb*ClpP1P2 (resolution 2.7 Å) is 6IW7.

### DSC

DSC-based analysis of the thermal denaturation of proteins provides an approach for measuring protein–ligand interactions[40]. Samples for DSC were prepared following the operating manual of the instrument. *Mtb*ClpP1P2 was dissolved to a final concentration of 0.5 mg/mL. The molar ratio between *Mtb*ClpP1P2 and compounds was 10:1 in reaction cells. DSC measurements were performed using a VP-DSC Micro Calorimeter (Microcal, USA) at a scan rate of 0.5–2°C/min in the temperature range 10–110°C. Six-pair blank cells with buffer (50 mM phosphate buffer, pH 7.6, 300 mM KCl, 5 mM MgCl_2_ and 5% glycerol, with the same volume of DMSO) were prepared to obtain instrumental baselines, which were systematically subtracted from the sample experimental thermograms. A thermal transition curve was obtained from a plot of heat capacity against temperature.

### Mutagenesis

Thirty-one mutations were introduced by site-directed mutagenesis, primers are listed in Appendix Table S3. The PCR products were extracted with PCR purification kits (Takara 9761) and ligated using a Blunting Kination Ligation Kit (Takara 6217). The mutated sequences of the ClpP1 gene were confirmed by DNA sequencing (Tsingke). Variant ClpP1 proteins were expressed in *Escherichia coli* BL21 (DE3) for enzymatic cleavage assays. The IC_50_ values of cediranib toward *Mtb*ClpP1P2 mutants were determined in the same way as that for wild-type *Mtb*ClpP1P2.

### Molecular Dynamics Simulations

The molecular dynamics simulations were performed in AMBER 14 software package[41]. The initial coordinates for molecular dynamics simulation were firstly prepared through structural inspection and optimization in Schrödinger software suite[42]. In the tleap module of AMBER, *Mtb*ClpP1P2 dimer was solvated in a rectangular water box of TIP3P and neutralized with Na+ ions. The periodic boundary conditions were setup with all the solvents 10 Å away from the solutes. The protein were parameterized using the AMBER FF99SB force field[43] and the ligands were parameterized using the GAFF force filed[44]. Energy minimization was performed firstly to remove the local atomic collision in the systems and the combination of the descent steepest with conjugated gradient method was adopted in the process. In the NVT ensemble, the systems were heated from 0 to 310 K gradually and the solutes were restrained with harmonic force constant 5 kcal/mol/Å2 simultaneously. Five equilibration stages were performed to adjust the solvents density. Finally, 500ns conventional molecular dynamics (cMD) simulations were performed in the NPT ensemble, with coordinates saved every 5 ps throughout the entire process.

### Data analysis for MD simulation

Root-mean-square deviation (RMSD) for the proteins and ligands during the simulations were calculated using the CPPTRAJ module[45] of AmberTools 13 package[41] to characterize the conformational change of the proteins and ligands. Standardized secondary structures assignment analysis were calculated with the DSSP algorithm using the AmberTools 13 package, which can characterize the propensities of secondary structures for each residue during the simulations. Then the heatmap for DSSP analysis was plotted using the MATLAB package (MathWorks, USA).Principal Component Analysis (PCA) analysis on the basis of covariance matrix was carried out using the program Carma[46] to address the collective motions of *Mtb*ClpP1P2. A two-dimensional representation of Free Energy Landscape (FEL) was built based on the PCA analysis, and two dominant components of PC1 and PC2 were selected as the reaction coordinates. The FEL along the reaction coordinates could be calculated using the following equation[47]:

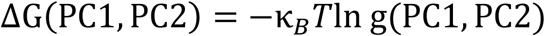

where T and κB represent the temperature of MD simulations and the Boltzmann constant, respectively. The g(PC1,PC2) is the normalized joint probability. Then the structures for the lowest energy in FEL were extracted as the representative conformation of each energy basin. These representative structures were superimposed to the initial crystal structure for comparison.

## Acknowledgements

We would like to thank Professor Jikui Song for generously sharing the plasmids used to generate ClpPs proteins. X-ray diffraction image collection, analysis and computation work were performed using the workstations at Shanghai Synchrotron Radiation Facility.

## Author contributions

**Conceptualization:** Youfu Luo.

**Data curation:**Yang Yang, Yibo Zhu and Huanxiang Liu.

**Formal analysis:** Yang Yang, Yibo Zhu, Tao Yang, Tao Li, Yuan Ju, Yingjie Song, Huanxiang Liu, Jun He, Rui Bao and Youfu Luo.

**Funding acquisition:** Rui Bao and Youfu Luo.

**Investigation:** Yang Yang, Yibo Zhu, Tao Yang, Tao Li, Yuan Ju, Yingjie Song, Jun He and Huanxiang Liu.

**Resources:** Rui Bao and Youfu Luo.

**Supervision:** Tao Yang, Rui Bao and Youfu Luo.

**Validation:** Yang Yang, Yibo Zhu, Tao Yang, Rui Bao and Youfu Luo.

**Visualization:** Yang Yang, Yibo Zhu and Rui Bao.

**Writing-original draft:** Yang Yang, Yibo Zhu, Tao Yang, Rui Bao and Youfu Luo.

**Writing-review & editing:** Yang Yang, Rui Bao and Youfu Luo.

## Competing interests

The authors declare no competing interests.

## Supporting Information

**S1 Table.**
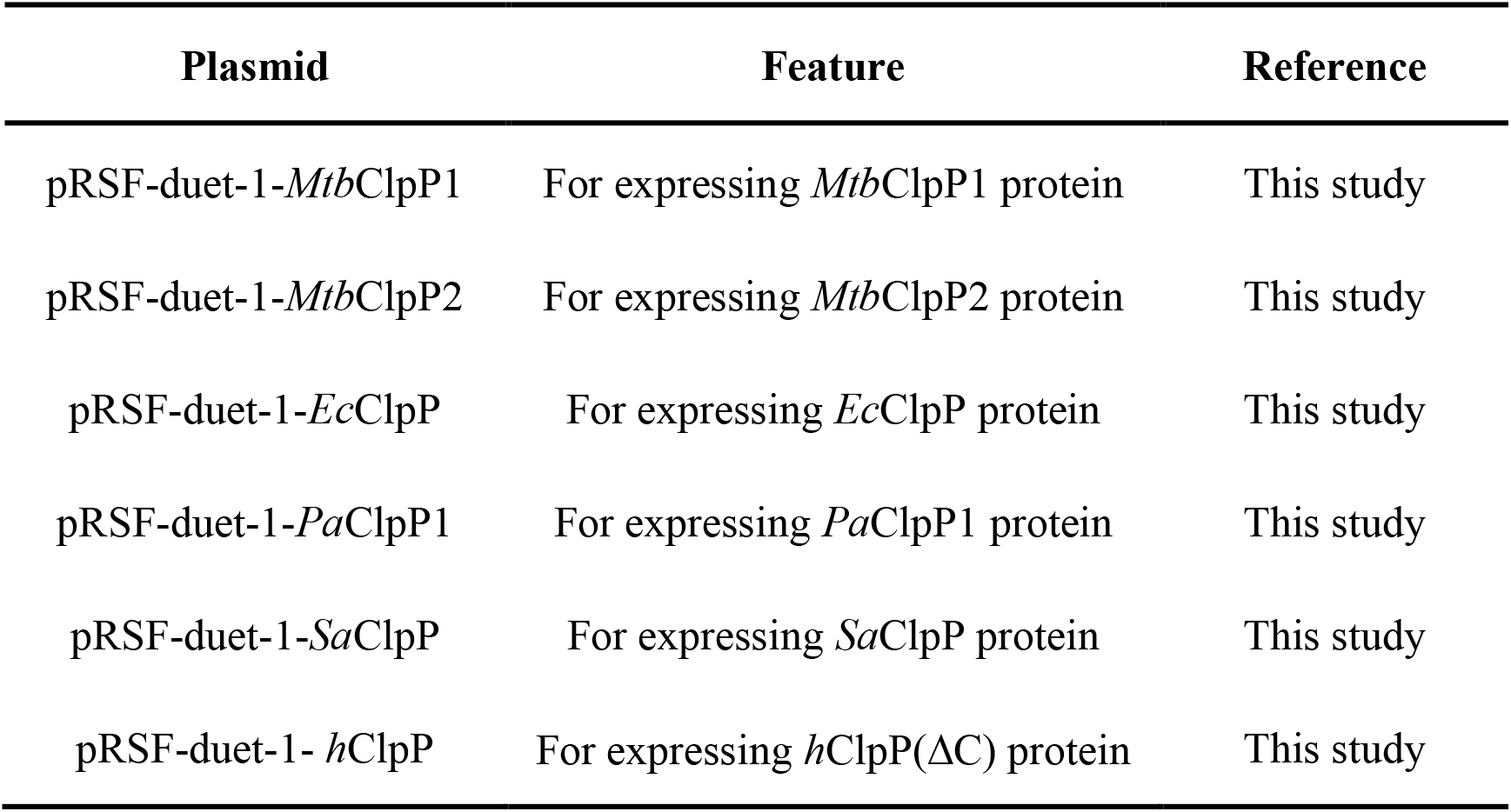
Plasmids used in this study.

**S2 Table.**
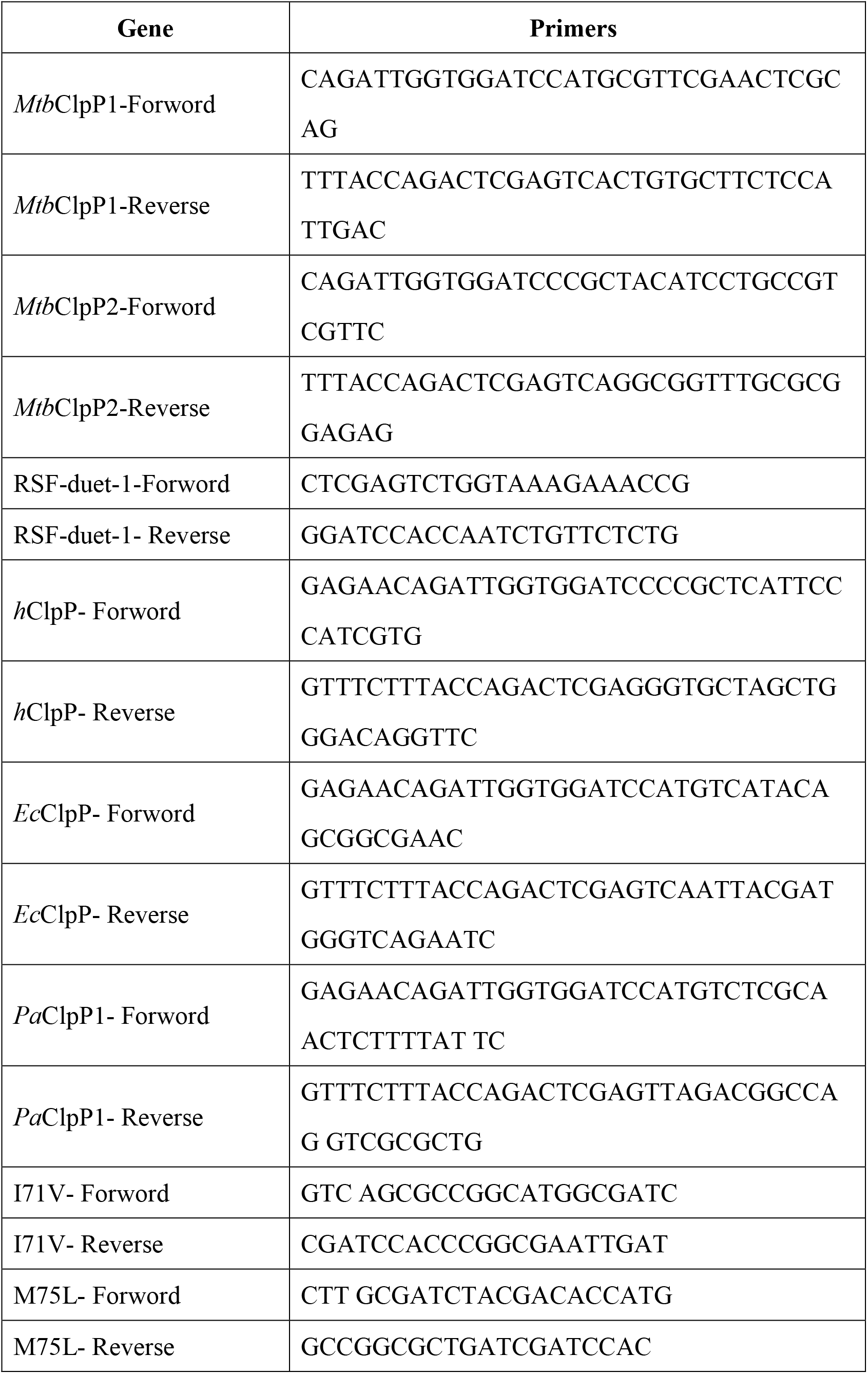
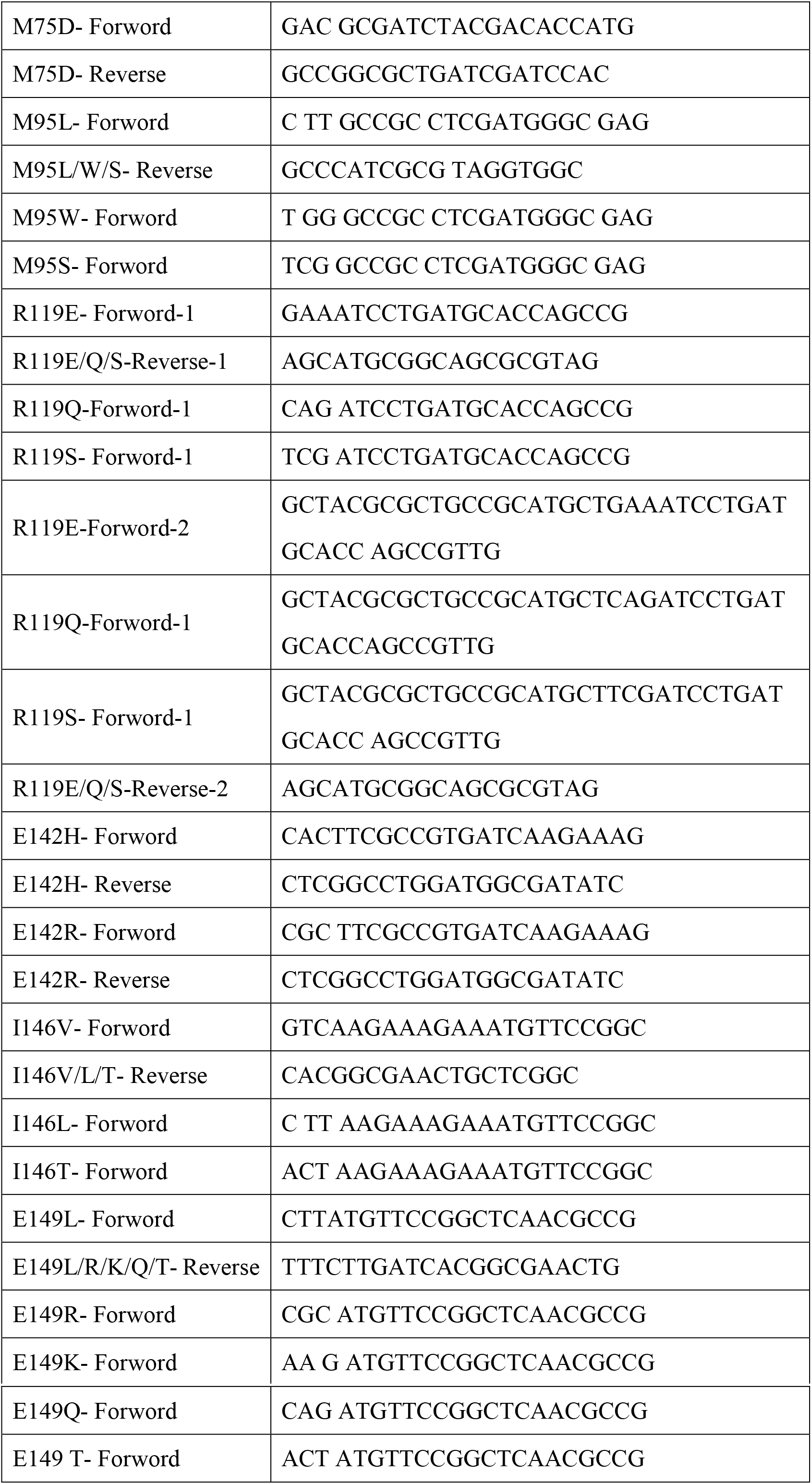
Primers used in this study.

**S3 Table.**
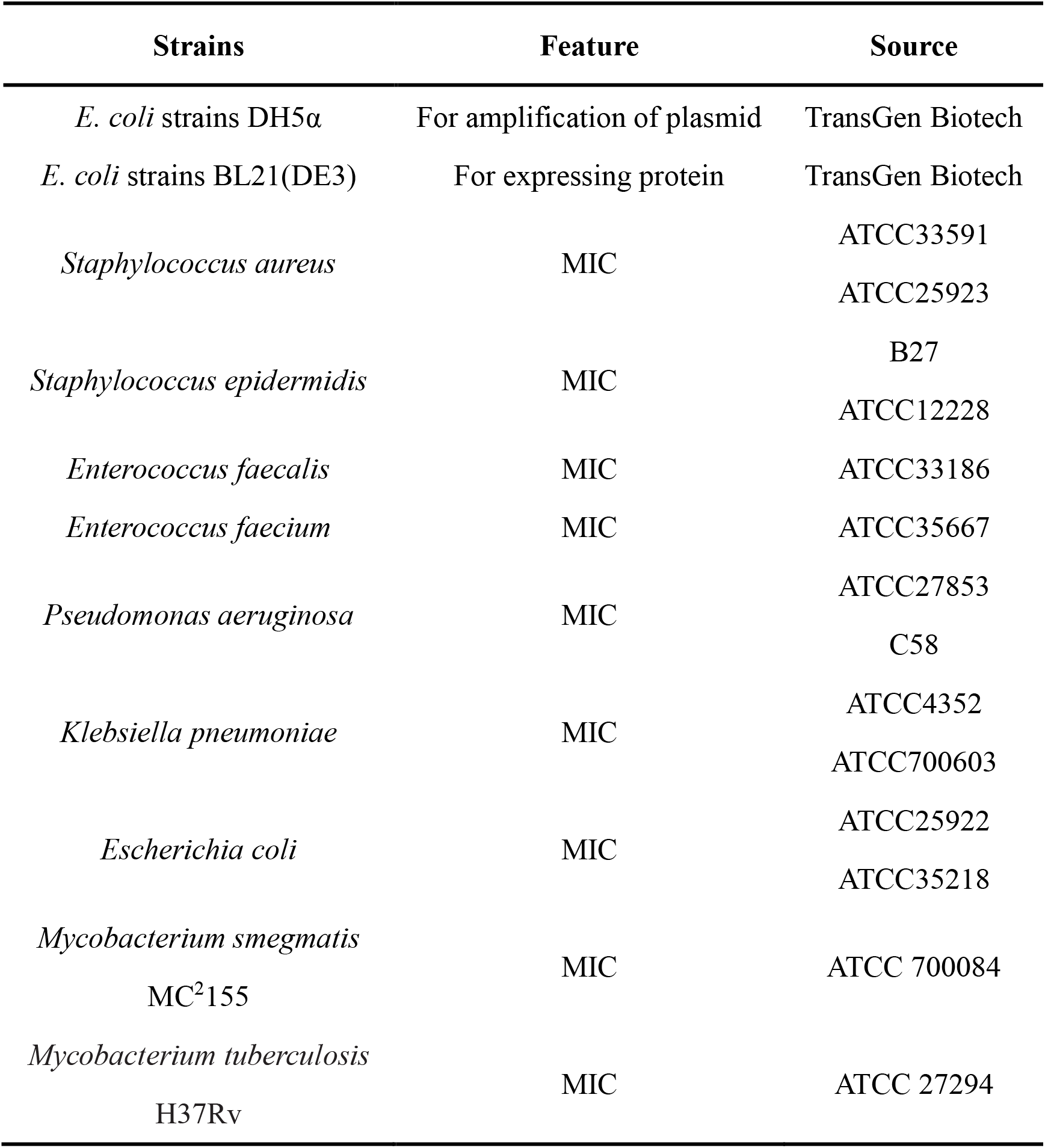
Strains used in this study.

**S1 Fig.**
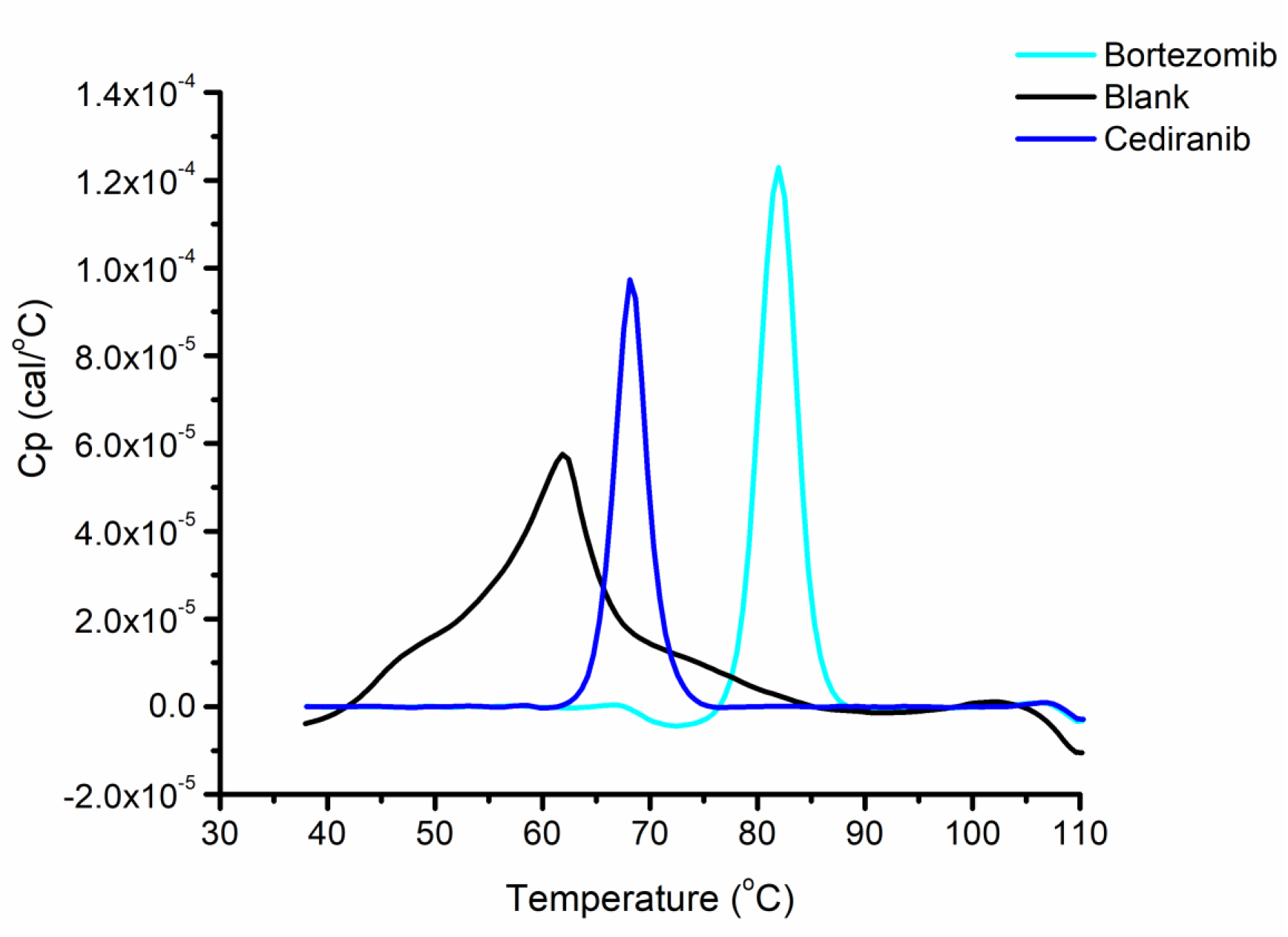
Differential scanning calorimetry analysis of the binding between cediranib and *Mtb*ClpP1P2. The concentrations of bortezomib and cediranib were 100 μM.

**S2 Fig.**
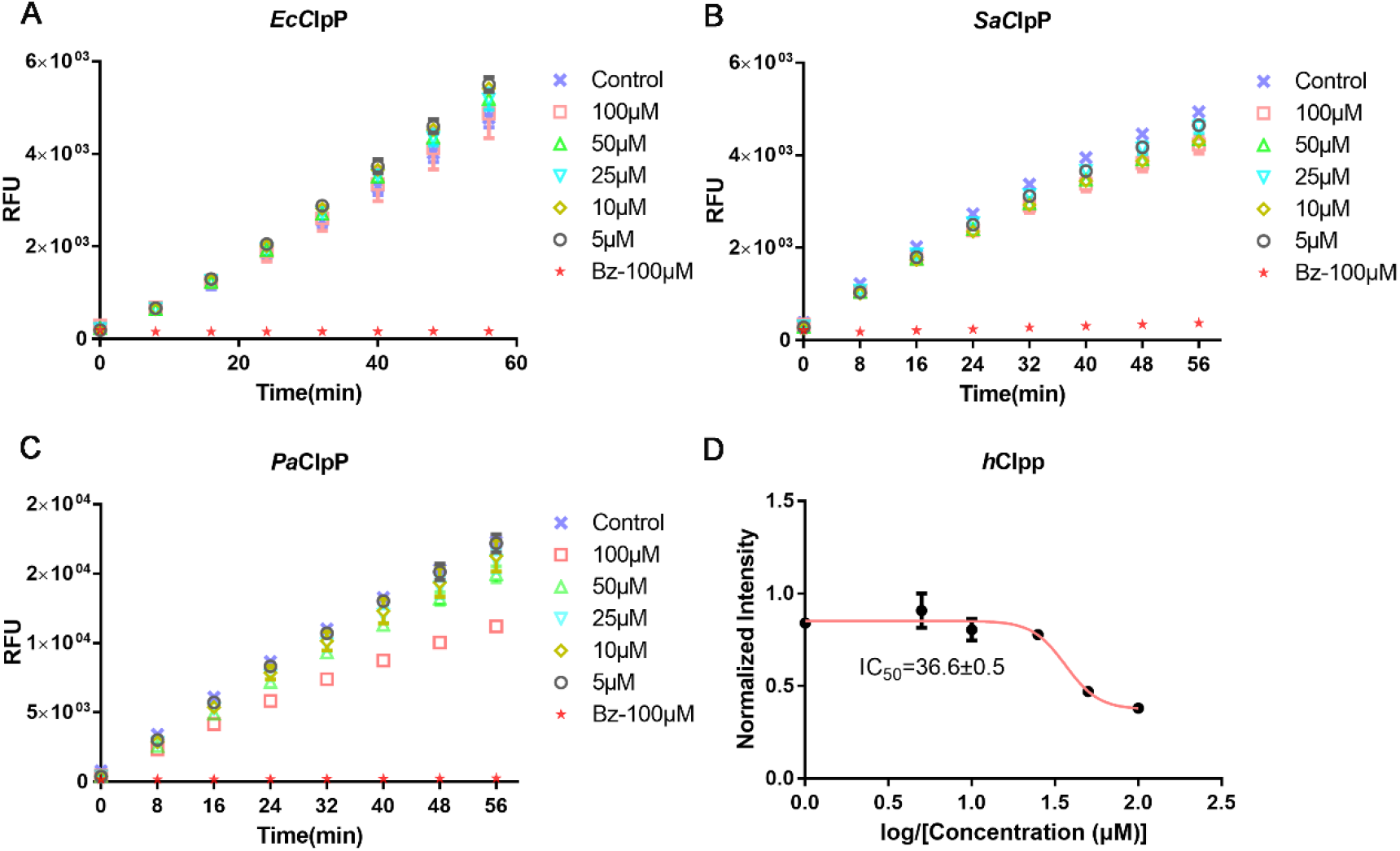
Inhibition of cediranib aganist *Ec*ClpP, *Sa*ClpP, *Pa*ClpP and *h*ClpP. (A) Inhibition of cediranib on *Ec*ClpP. Concentration of *Ec*ClpP was 0.5μM. (B) Inhibition of cediranib on *Sa*ClpP. Concentration of *Sa*ClpP was 2.5 μM. (C) Inhibition of cediranib on *Pa*ClpP. Concentration of *Pa*ClpP was 0.5 μM. (D) IC_50_ value of cediranib aganist *h*ClpP. Concentration of *h*ClpP was 2.5μM.

**S3 Fig.**
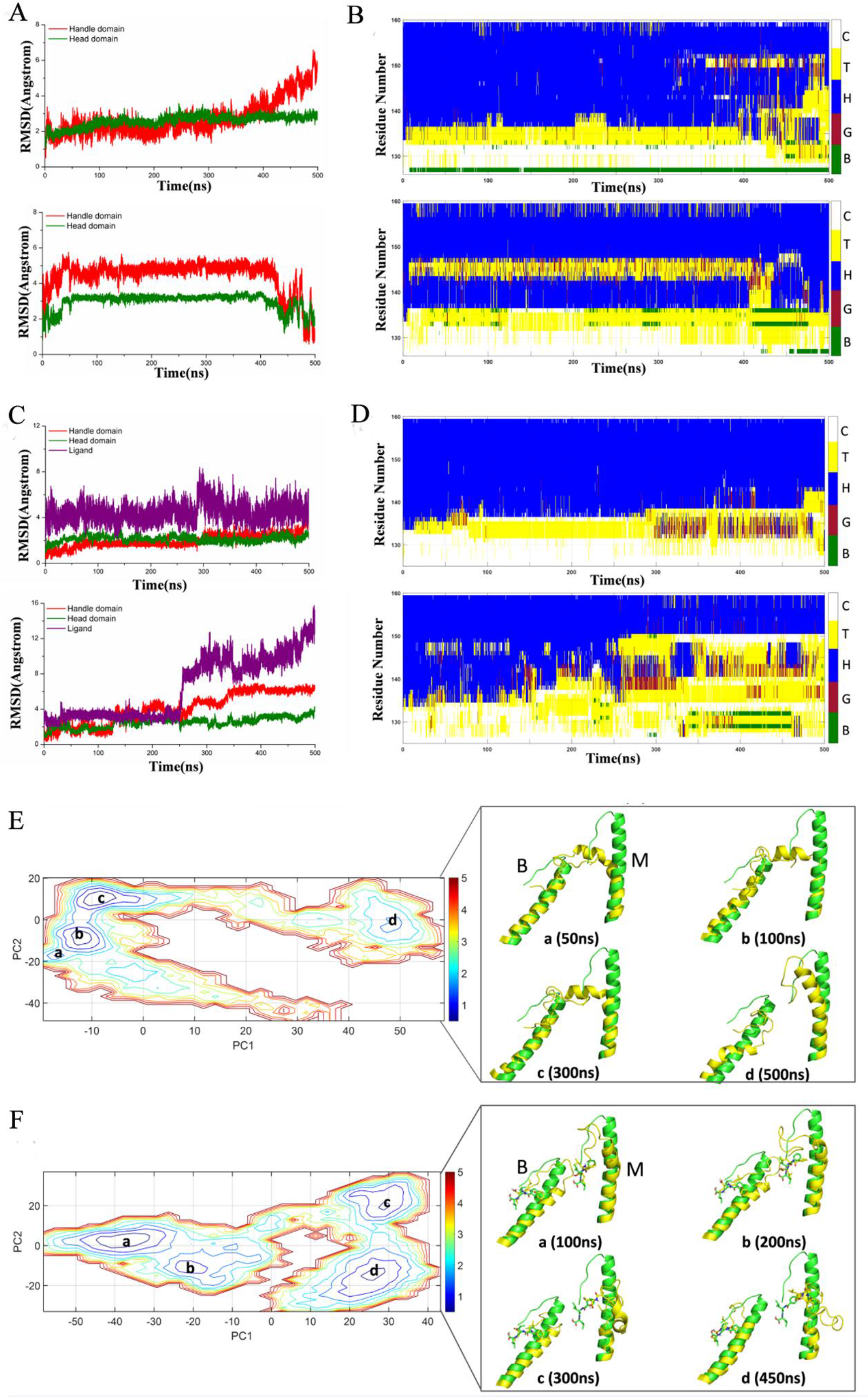
Conformational changes of *Mtb*ClpP1 dimer. (A,C). Cα RMSD values of the head domain and handle domain in *Mtb*ClpP1 dimer versus simulation time without (A)or with cediranib(C). The RMSD values of the head domain and handle domain are shown in green and red, respectively. (B,D). Secondary structures as a function of time for *Mtb*ClpP1 dimer without (B)or with cediranib(D) in trajectory as calculated using DSSP. The structures were analyzed every 100 ps. (E,F). Left, energy landscape for the conformational transition of *Mtb*ClpP1 dimer without (E)or with cediranib(F). Reaction coordinates were defined according to PC1 and PC2 obtained from PCA. Right, snapshot structures of α5 from *Mtb*ClpP1 dimer without (E)or with cediranib(F) extracted from the trajectory.

